# Endothelial TIE1 restricts angiogenic sprouting to coordinate vein assembly in synergy with its homologue TIE2

**DOI:** 10.1101/2022.08.05.502976

**Authors:** Xudong Cao, Taotao Li, Beibei Xu, Kai Ding, Weimin Li, Bin Shen, Man Chu, Dengwen Zhu, Li Rui, Zhi Shang, Xiao Li, Yinyin Wang, Shuyu Zheng, Kari Alitalo, Ganqiang Liu, Jing Tang, Yoshiaki Kubota, Yulong He

**Author notes:** Correspondence should be addressed to Dr. Yulong He. Equal contribution (Xudong Cao, Taotao Li, Beibei Xu).

## Abstract

**Objective:** Vascular growth followed by vessel specification is crucial for the establishment of a hierarchical blood vascular network. We have here investigated mechanisms underlying venogenesis, particularly the molecular control over venous fate acquisition during vascular development.

**Approach and Results:** We analyzed the function of TIE1 as well as its synergy with TIE2 in the regulation of vein formation by employing genetic mouse models targeting *Tie1* and *Tek*. Cardinal vein growth appeared normal in TIE1 deficient mice, whereas TIE2 deficiency altered the identity of cardinal vein endothelial cells with the aberrant expression of DLL4. Interestingly, the parallel growth of murine cutaneous veins along with arteries, which was initiated at approximately embryonic day 13.5, was retarded in mice lack of TIE1. *Tie1* deletion disrupted also venous integrity, displaying increased sprouting angiogenesis and vascular bleeding. Abnormal venous sprouts with defective arteriovenous alignment were also observed in the mesenteries of *Tie1* deleted mice. Mechanistically, TIE1 deficiency resulted in the decreased expression of venous regulators including TIE2 while angiogenic regulators were upregulated. The alteration of TIE2 level by TIE1 insufficiency was further confirmed by the siRNA-mediated knockdown of *Tie1* in cultured endothelial cells. Additionally, combining the endothelial deletion of *Tie1* with one null allele of *Tek* resulted in a progressive increase of vein-associated angiogenesis leading to the formation of vascular tufts in retinas, whereas the loss of *Tie1* alone produced only a relatively mild venous defect.

**Conclusions:** Findings from this study imply that TIE1 and TIE2 act in a synergistic manner to restrict sprouting angiogenesis during vein formation.

## Introduction

Specification of vascular endothelial cell (EC) identity, together with the parallel formation of arteries and veins, are crucial events for the construction of vascular network in development. While we have gained a better understanding about the molecular players for arteriogenesis, mechanisms underlying vein development are less well characterized. COUP-TFII, a transcription factor expressed in venous ECs, has been shown to regulate venous identity via the inhibition of NOTCH-mediated signals [1]. AKT activation was shown to inhibit RAF1-ERK1/2 signaling in ECs to favor venous specification [2]. Recently the upstream regulators have also been identified. The angiopoietin (ANGPT) receptor TIE2 was demonstrated to play a crucial role in the venous specification and maintenance via its downstream PI3K/AKT-mediated stabilization of COUP-TFII [3]. Mutations with angiopoietin receptor TIE2 led to venous malformation [4]. Consistently, the specific deletion of cardiomyocyte *Angpt1* was shown to disrupt coronary vein formation in the developing heart [5]. Combined insufficiency of TIE2 ligands ANGPT1 and ANGPT2 in mice disrupted the formation of sclera venous sinus (Schlemm’s canal), a type of vessel with also lymphatic characteristics [6, 7].

TIE1 is a member of the receptor tyrosine kinase family with a high degree of homology with TIE2. Genetic studies have revealed that angiopoietins and TIE receptors are differentially required during blood vascular and lymphatic development. Attenuation of ANGPT1 and TIE2-mediated signals disrupted vein formation in addition to other vascular phenotypes [3, 5, 8-10]. *Tie1* and *Angpt2* null mice displayed abnormal lymphatic formation and also blood vascular defects [11-18]. In spite of the important roles of TIE1 in vascular development, the underlying mechanism of its functions remains incompletely understood. On the one hand, previous studies showed that TIE1 could heterodimerize with TIE2 and exert an inhibitory role in TIE2 signaling [19-21]. TIE1 expression was also suggested to negatively regulate TIE2 presentation at the cell surface in sprouting endothelial tip cells [17]. On the other hand, ANGPT1 could induce TIE1 phosphorylation when co-expressed with TIE2 in cultured cells [22]. TIE1 and TIE2 heteromeric complexes in endothelial cell–cell junctions were shown to be required for TIE2 activation [16, 23, 24].

By employing genetic mouse models targeting *Tie1, Tek* (encoding TIE2) or both, we have explored functions of TIE1 and its relationship with TIE2 in the process of venogenesis. We found in this study that vein formation was retarded in TIE1 deficient mice, which showed an increase of vein-associated angiogenic sprouting during embryogenesis as well as in the postnatal retinal vascular development. Loss of TIE1 led to the decrease of TIE2 on the levels of messenger RNA and protein. Combining the endothelial deletion of *Tie1* plus one null allele of *Tek* led to the formation of retinal vein-associated vascular tufts similar to that observed in the endothelial *Tek* deleted mice. Findings from this study imply that TIE1 acts in synergy with TIE2 to restrict angiogenesis for venous assembly during the establishment of a hierarchical vascular network.

## Material and Methods

### Mouse models

All animal experiments were performed in accordance with the institutional guidelines of Soochow University Animal Center. Two genetically modified mouse models targeting *Tie1* gene were employed in this study. One mouse line targets TIE1 intracellular kinase domain (ICD), with exon 15 and exon 16 floxed, *Tie1ICD*^*Flox/Flox*^ [12]. The other line is a knock-out first mouse model established from EUCOMM embryonic stem cells (EPD0735-3B07) targeting *Tie1* gene (*Tie1*^*tm1a/tm1a*^), in which targeting cassette is recombined downstream of exon 7 (with exon 8 floxed). *Tek* knockout mouse model was generated as previously reported [3] and was crossbred with *Tie1*^Δ*ICD/*Δ*ICD*^ mouse model to obtain T*ie1ICD* and *Tek* double knockout mice (*Tie1*^*+/*^Δ*ICD;Tek*^*+/-*^). To generate mice with endothelial cell specific gene deletion mouse models, we employed the *Cdh5-Cre*^*ERT2*^ mouse line [25]. In all the phenotype analysis, littermates were used as controls. Mice were bred in SPF experimental animal facilities. For the genotyping of *Tie1*^*tm1a*^ knockout allele, the primers used were forward primer (5’- GCATGAAACTTCGCAAGCCA -3’) and reverse primer (5’- CTCTGCTGTGGTCCTGTCTG -3’), to amplify a 326 bp fragment for the wildtype allele and a 387 bp for the knockout allele. For the genotyping of *Tie1ICD* and *Tek* mouse lines, the primers used were as previously described [12].

### Induced gene deletion

Induction of gene deletion was performed as previously described by tamoxifen treatment [3]. Briefly, new-born pups from the knockout (*Tie1ICD*^*ECKO/-*^ and *Tie1ICD*^*ECKO/-*^*;Tek*^*+/-*^) and control mice were treated by the intragastric injection from postnatal day 1 for 4 days. Tissues were collected for analysis at postnatal day 7-21. The retina dissection was according to the protocol by Pitulescu et al. [26]. Retinal vascularization index was quantified as the ratio of vascularized area to total retinal area as previously published {Chu, 2016 #8258}.

### RNA sequencing analysis

Skin tissues were harvested from *Tie1* knockout and control mice (E17.5) and kept frozen in liquid nitrogen. Total RNA was extracted from the tissues using Trizol (Invitrogen) according to the manufacturer’s instruction. RNA was qualified and quantified using a Nano Drop and Agilent 2100 bioanalyzer (Thermo Fisher Scientific). Oligo(dT)-attached magnetic beads were used to purify mRNA. Purified mRNA was fragmented, converted into cDNA and amplified by PCR, and finally made into RNAseq libraries. Single end 50 bases reads were generated with a target depth of 20 million reads on BGIseq500 platform (BGI-Shenzhen, China). The sequencing data was filtered with SOAPnuke (v1.5.2), and clean reads were obtained and stored in FASTQ format. The clean reads were mapped to the reference genome using HISAT2 (v2.0.4). Bowtie2 (v2.2.5) was applied to align the clean reads to the reference coding gene set. Gene expression level was calculated by RSEM (v1.2.12), and differential expression analysis was performed using the DESeq2 (v1.4.5) with Q value ≤ 0.05 [27-31]. Gene enrichment analysis was performed using GSEA R package clusterProfiler (v3.14.3) [32-34].

### RT-PCR

Total RNA from the lung and skin tissues of *Tie1*^Δ*ICD/*Δ*ICD*^ and control embryos (E15.5) were extracted by TRIzol following the manufacturer’s protocol. cDNA was synthesized using a reverse transcriptional reaction kit (RevertAid First Strand cDNA Synthesis Kit, Thermo Scientific). Real-time quantitative PCR was performed using a SYBR Premix Ex Taq kit (Takara RR420A) in the Applied Biosystems 7500 Real-Time PCR System. Primers of qRT-PCR are as follows: *Gapdh*: 5’-GGTGAAGGTCGGTGTGAACG-3’,5’-CTCGCTCCTGGAAGATGGTG-3’; *Tie1*: 5’-GCTGTGGTAGGTTCCGTCTC-3’, 5’-AAGGTCCCTGAGCTGAACTG-3’; *Tek*: 5’-GATTTTGGATTGTCCCGAGGTCAAG-3’, 5’-CACCAATATCTGGGCAAATGATGG-3’; *Aplnr*: 5’-CAGTCTGAATGCGACTACGC-3’, 5’-CCATGACAGGCACAGCTAGA-3’; *Efnb2*: 5’-TGTTGGGGACTTTTGATGGT-3’, 5’-GTCCACTTTGGGGCAAATAA-3’; *Notch1*: 5’-TGTTGTGCTCCTGAAGAACG-3’, 5’-TCCATGTGATCCGTGATGTC-3’; *Dll4*: 5’-TGCCTGGGAAGTATCCTCAC-3’, 5’-GTGGCAATCACACACTCGTT-3’. The transcripts of venous and arterial markers were normalized against *Gapdh*, and the relative expression level of every gene in the *Tie1*^Δ*ICD/*Δ*ICD*^ mice was normalized against that of littermate control mice.

### Cell culture and siRNA transfection

Human umbilical vein endothelial cells (HUVECs) were cultured in endothelial cell medium (ScienCell Research Laboratories #1001). To knock down *Tie1* expression in HUVECs, cells were transfected with siRNA targeting human *Tie1* (s14141 or s14142; Invitrogen) using Lipofectamine RNAiMax (Invitrogen), and siRNA negative control duplexes (12935300; Invitrogen) were used as a control.

### Western blotting analysis

Lung and skin tissues of embryos were harvested, snap-frozen in liquid nitrogen and stored at -80 °C freezer. Tissues or cells were lysed in NP-40 lysis buffer (Beyotime P0013F) supplemented with protease inhibitor cocktail (complete Mini, Roche 04693124001), phosphatase inhibitor cocktail (PhosSTOP, Roche 04906837001), 10 mM NaF and 1 mM PMSF. Protein concentration was determined using the BCA protein assay kit (PIERCE), and equal amounts of protein were used for analysis. The following primary antibodies were used in this study, including rabbit polyclonal anti-TIE2 (Santa Cruz sc-324), goat polyclonal anti-TIE2 (R&D Systems AF762), goat polyclonal anti-TIE1 (R&D Systems AF619), rabbit polyclonal anti-AKT (Cell Signaling Technology #9272), rabbit monoclonal anti-Phospho-AKT (Ser473, Cell Signaling Technology #4060), mouse monoclonal anti-COUP-TFII (R&D Systems #PP-H7147-00), mouse monoclonal to beta-Actin (Santa Cruz sc-47778).

### Immunostaining

For the embryo studies, female mice were mated in the late afternoon and vaginal plugs were checked in the morning of the following day. The embryonic stages are estimated considering midday of the day on which the vaginal plug is present as embryonic day 0.5 (E0.5). Yolk sacs or tail tips from embryos were collected for genotyping. For immunostaining, tissues were harvested and processed as previously described [35]. Briefly, the tissues were fixed in 4% paraformaldehyde, blocked with 3% (w/v) skim milk in PBS-TX (0.3% Triton X-100), and incubated with primary antibodies overnight at 4°C. The antibodies used were: rat anti-mouse PECAM1 (BD 553370), 660-mouse-anti-mouse αSMA (eBioscience 50-9760), Cy3-mouse-anti-mouse aSMA (Sigma C6198), mouse anti-mouse αSMA (Sigma A2547), goat-anti-human TIE1 (R&D AF619), goat-anti-mouse TIE2 (R&D AF762), goat-anti-mouse DLL4 (R&D AF1389), goat-anti-mouse EphB4 (R&D AF446), rat anti-mouse endomucin (eBioscience 14-5851), rabbit anti-all NG2 (Chemicon AB5320). Appropriate Alexa 488, Alexa594 (Invitrogen) conjugated secondary antibodies were used. All fluorescently labeled samples were mounted and analyzed with a confocal microscope (Olympus Flueview 1000), or Leica MZ16F fluorescent dissection microscope.

### Statistical analysis

For the two-group comparison, the unpaired *t* test was performed if data passed the D’Agostino-Pearson normality test, or the unpaired nonparametric Mann-Whitney test was applied using GraphPad Prism 7. All statistical tests were 2-sided.

## Results

### Differential requirement of TIE1 and TIE2 in cardinal vein specification

To investigate if TIE receptors are required for the specification of cardinal veins, we analyzed the *Tie1* and *Tek* mutant mice during early embryogenesis. As shown by immunofluorescent staining of the pan-endothelial marker PECAM-1, plus TIE1 or TIE2 in Fig. 1A (embryonic day 8.75, E8.75), both TIE receptors were expressed in the dorsal aortas (DA) and anterior cardinal veins (ACV). To further characterize their functions in the cardinal vein formation, we deleted *Tie1, Tek* or both as previously reported [3, 12]. We found that the cardinal veins as well as dorsal aortas (7 to 14-somite stages, E8.75) appeared normal in *Tie1* deleted mice (Fig. 1B), with DLL4 expression in the dorsal aortas but not in cardinal veins. However, TIE2 deficiency led to the upregulation of DLL4 in anterior cardinal veins (Fig. 1C). This result is consistent with the previous observation that TIE2 is required for the specification of venous identity [3]. The double deletion of both *Tie1* and *Tek* (*Tie1*^Δ*ICD/*Δ*ICD*^*;Tek*^*-/-*^) resulted in the increased DLL4 expression in ACV and posterior cardinal veins (PCV) as shown in Fig. 1D. Furthermore, the double-deleted embryos showed an obvious decrease of blood vessel density in head regions (asterisk in Fig. 1E).

**Fig. 1.**
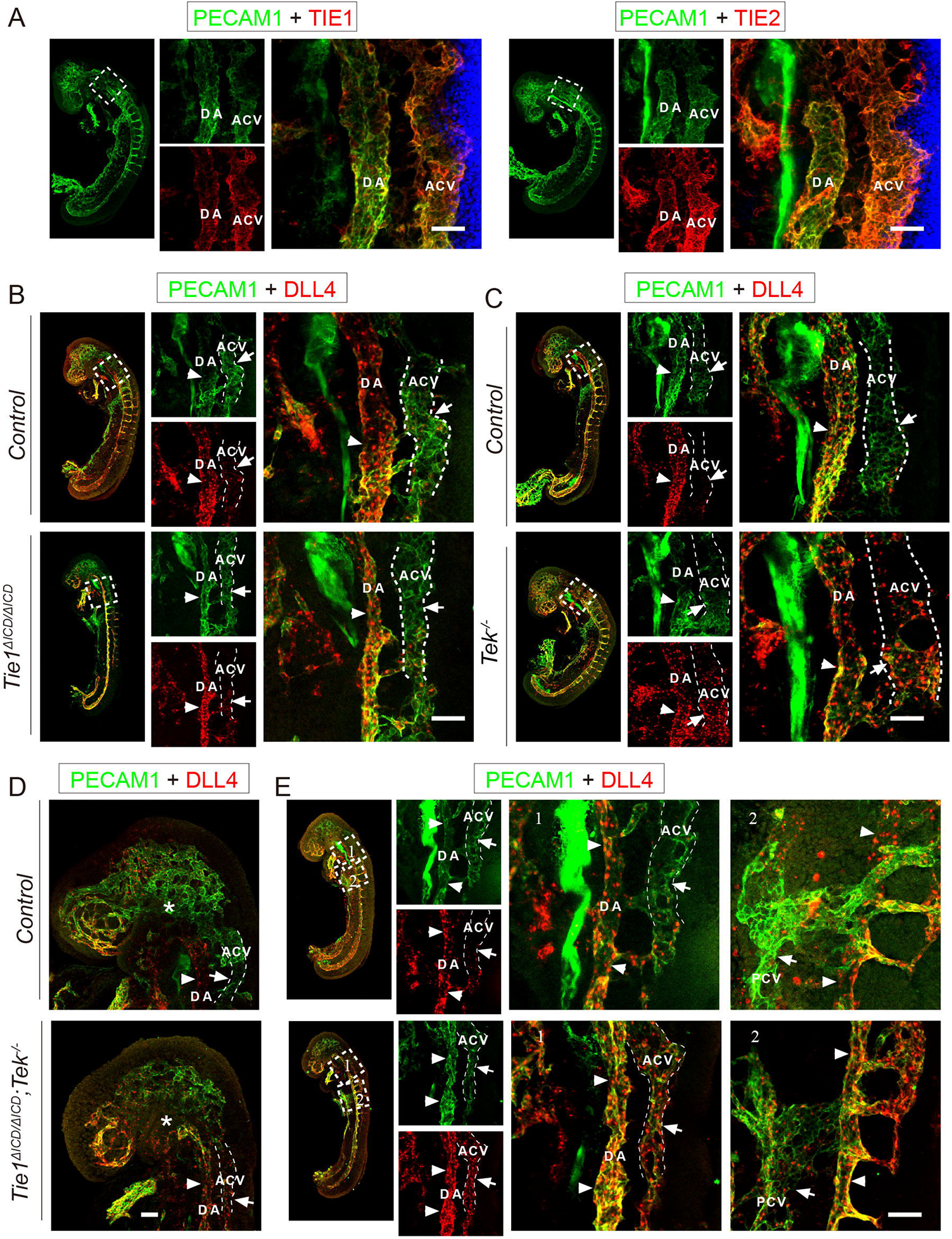
Differential requirement of TIE1 and TIE2 in cardinal vein specification at early embryogenesis. **A**. Analysis of cardinal veins and aortas (E8.75) by the wholemount immunostaining for PECAM1 (green) and TIE1 or TIE2 (red). **B-E**. *Tie1* deletion did not affect cardinal vein formation (B), while *Tek* deletion altered the endothelial cell identity of cardinal veins (C-E; E8.5-9.0, 7 to 14-somite stages). Expression of arterial marker DLL4 (red, green for PECAM1) was detected in cardinal veins. Vein formation was disrupted in *Tek* null (*Tek*^*-/-*^, C) or *Tie1/Tek* double knockout mice (*Tie1*^Δ*ICD/*Δ*ICD*^;*Tek*^*-/-*^, D, E). Asterisk in D points to DLL4 negative vessels, which are absent in *Tie1/Tek* double knockout mice. Arrows point to anterior cardinal veins and arrowheads to dorsal aortas. DA, dorsal aorta; ACV, anterior cardinal vein; PCV, posterior cardinal vein. Scale bar: 50 μm in A-E.

### Disruption of cutaneous vein formation and alignment with arteries after *Tie1* deletion

TIE2 is known to be more abundant in veins than arteries during early embryogenesis [3]. Also, TIE1 was detected mainly in veins in head (Fig. 2A) and somite regions at E10.5 (Fig. 2B). Although TIE2 deficiency disrupted the vein formation in head and somite regions as observed at E9.5 [3], lack of TIE1 did not have an obvious effect on vein formation (Fig. 2A and B, E10.5), or the recruitment of vascular mural cells in head regions (Supplemental Fig. 1A). However, the vein formation was disrupted in the skin of *Tie1* mutant mice in comparison with the heterozygous and wildtype controls (Fig. 2C, and Supplemental Fig. 1C), in addition to an increase of non-vascularized regions in the dorsal skin (Supplemental Fig. 1B) and the defective lymphatic development (Supplemental Fig. 2). Small arteries but no veins were detected in regions close to the midline of the dorsal skin in the *Tie1*^Δ*ICD/*Δ*ICD*^ mice (Fig. 2D). Blood vessels in these regions also showed more angiogenic sprouting and less pericyte coverage in the *Tie1* mutants in comparison with the controls (Supplemental Fig. 1D). Interestingly, the veins were detected at the upper part of the dorsal cutaneous vascular network close to the axilla region (E15.5, Fig. 2D-E), but they were not properly aligned with the arteries (Fig. 2E).

**Fig. 2.**
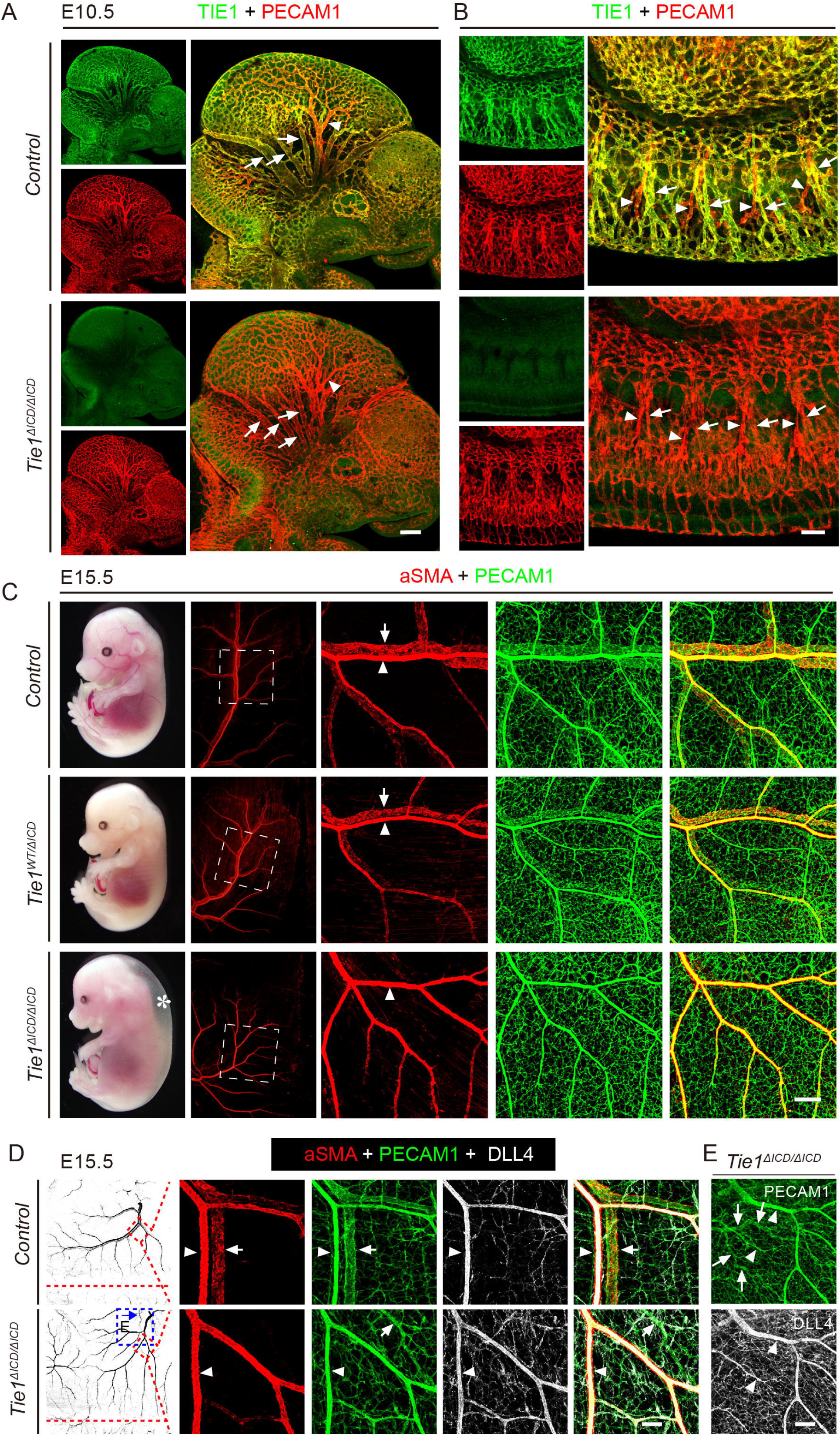
Defective vein formation after *Tie1* deletion. **A**,**B**. Visualization of blood vessels in head (A) and somite regions (B, E10.5) of *Tie1*^Δ*ICD/*Δ*ICD*^ and control mice by whole-mount immunostaining for PECAM1 (red) and TIE1 (green). Note that TIE1 is mainly expressed in veins (arrows) but with less expression in arteries in head and somite regions (arrowheads) at early stages of embryogenesis. **C**. Analysis of blood vessels in the skin of *Tie1* deleted and control mice (E15.5) by the immunostaining for PECAM-1 (green) and αSMA (red). Note that in comparison with the well-aligned veins and arteries in the skin of control mice, there were no veins but arteries detected in regions close to the midline of the dorsal skin in *Tie1* mutant mice. Subcutaneous edema was indicated by asterisk in *Tie1* mutants (C). **D, E**. Underdeveloped veins were accompanied by the increased staining of DLL4 in the skin of *Tie1*^Δ*ICD/*Δ*ICD*^ (E15.5) and the wildtype littermates were used as the control (PECAM1, green; αSMA, red; DLL4, white). Note that veins (DLL4 negative, arrows) were detected in the upper part of the dorsal cutaneous vascular network but not properly aligned with arteries in *Tie1* mutant mice. Arrows point to veins and arrowheads to arteries. Scale bar: 200 μm in A, C, E and 100 μm in B, D.

### Retardation of venous assembly upon TIE1 deficiency

As previously reported [12], the *Tie1*^Δ*ICD/*Δ*ICD*^ mutant mouse model was originally designed to investigate functions of TIE1 intracellular kinase domain in vascular development. However, because of the low level of expression of the truncated form of TIE1 lacking the intracellular domain (TIE1^ΔICD^), it is almost equivalent to a *Tie1* complete knockout mouse line. The lymphatic phenotype of *Tie1*^Δ*ICD/*Δ*ICD*^ mutants was similar to that observed in *Tie1* knockout mice [36]. To verify the role of TIE1 in the regulation of vein formation, we generated a new mouse line targeting *Tie1* gene (*Tie1*^*tm1a*^), which was a knockout first allele with *Tie1* exon 8 flanked by loxP sites (Supplemental Fig. 3A). *Tie1* deletion was confirmed by the PCR genotyping and western blotting analysis (Supplemental Fig. 3B, C). Sequential analysis of cutaneous vascular development at different stages of embryogenesis revealed that vein formation was initiated at approximately E13.5 close to the axilla region of dorsal skin, proceeding towards the midline (Fig. 3A). Veins aligned with the arteries were observed from E14.0 onwards in the control mice (E14.0-15.5; Fig. 3B-D). However, the development of veins was seriously retarded in *Tie1*^*tm1a/tm1a*^ mutant mice, being detected only at the upper part of dorsal cutaneous vascular network (Fig. 3B-D). As shown in Fig. 3E, the veins were not aligned properly with arteries unlike in the littermate control mice (Fig. 3E). Furthermore, DLL4 staining throughout the cutaneous vascular network was increased in the *Tie1* deleted mice in comparison with their littermate controls (Fig. 3C-D).

**Fig. 3.**
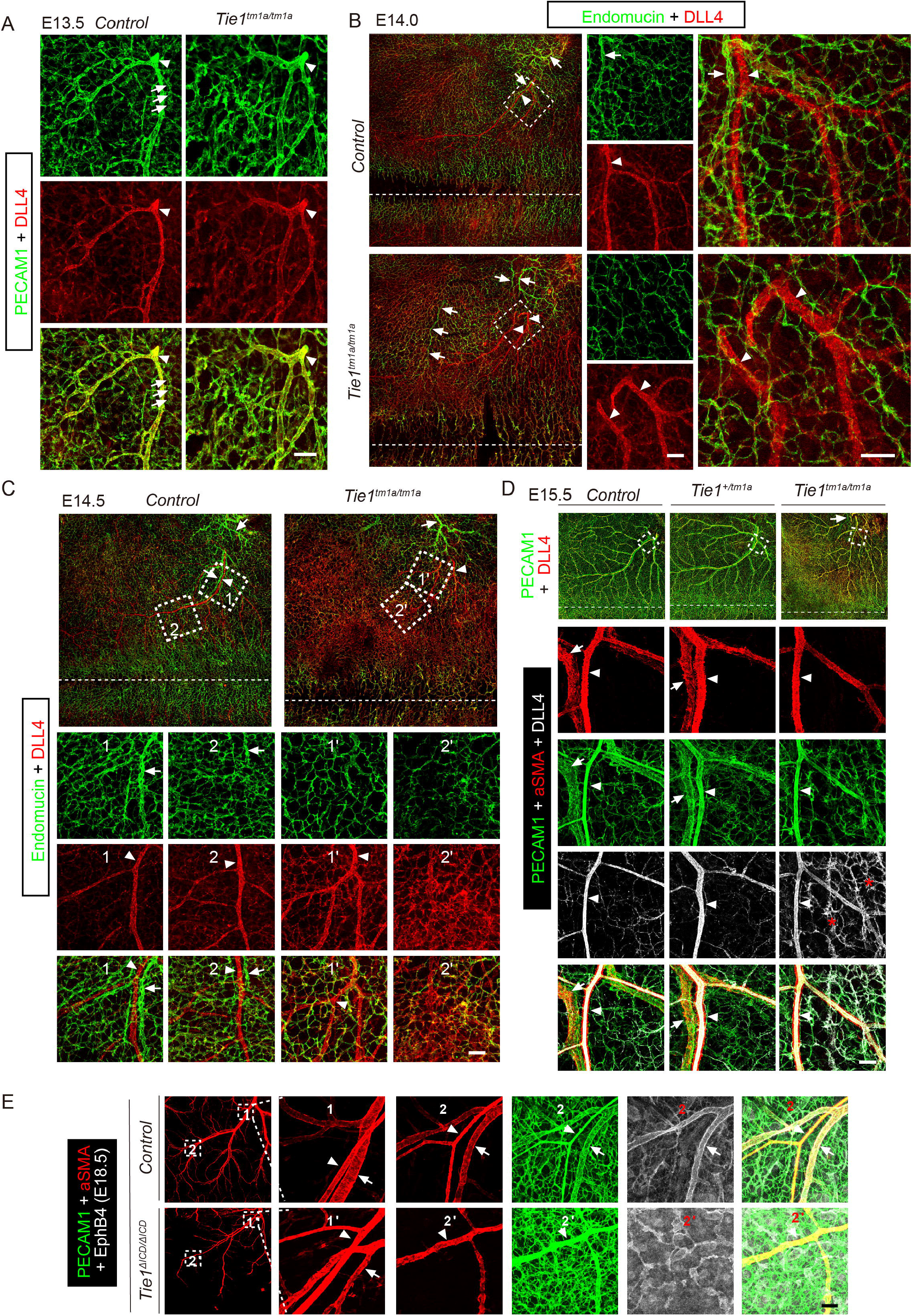
Requirement of TIE1 for the formation of cutaneous veins in alignment with arteries. **A**-**D**. Sequential analysis of vein formation in the skin of *Tie1*^*tm1a/tm1a*^ mice and littermate controls between E13.5-E15.5 (A, E13.5; B, E14.0; C, E14.5; D, E15.5) by whole-mount immunostaining for PECAM1 or Endomucin (green) and DLL4 (red). Note that arteries (DLL4 positive, arrowheads) were readily detected while the process of vein formation (PECAM1 positive but negative for DLL4) started at approximately E13.5 (A, arrows). Veins positive for Endomucin but negative for DLL4 were detected in the skin of *Tie1* mutant mice at the upper part of the vascular network near the axilla region of dorsal skin (B-D), but failed to grow along arteries (C, box 1’ and 2’), as compared with those of the control mice (C, box 1 and 2). **E**. Retardation and misalignment of veins in the skin of *Tie1*^Δ*ICD/*Δ*ICD*^ mice (E18.5). Consistent with the observation at E15.5, veins were detected at the upper part of the dorsal cutaneous vascular network in the skin of *Tie1* mutant mice (E18.5), but did not align properly with arteries as those in the littermate controls. An obvious increase of DLL4 staining in *Tie1* deleted mice was shown in C-D. Arrows point to veins and arrowheads point to arteries. Scale bar: 100 μm in A-E.

### Abnormal sprouting angiogenesis after the loss of TIE1

Consistent with the increased DLL4 expression in blood vessels of *Tie1* mutant mice, we found that lack of TIE1 was associated with abnormal vascular sprouting that impaired the venous integrity in the head regions (Fig. 4A). The delayed formation of veins was also observed in the mesentery of *Tie1*^*tm1a/tm1a*^ mice. The vascular structures positive for endomucin, expressed by venous but not arterial endothelial cells, were not properly formed in the mutant mice compared with the littermate controls (Fig. 4B, E13.5). In contrast to the mature mesentery veins aligned with arteries in the control mice (E17.5), mesentery veins of *Tie1* mutant mice still underwent active sprouting (Supplemental Fig. 3D). The immunostaining for VE-cadherin revealed that mesentery veins of *Tie1*^*tm1a/tm1a*^ mutants at this time point had irregular endothelial cell junctions and the venous endothelial cells were not properly elongated along the blood flow, whereas no obvious defects were observed in arteries (Fig. 4C). Similar venous abnormalities were observed in *Tie1*^Δ*ICD/*Δ*ICD*^ mice (Supplemental Fig. 4A-B).

**Fig. 4.**
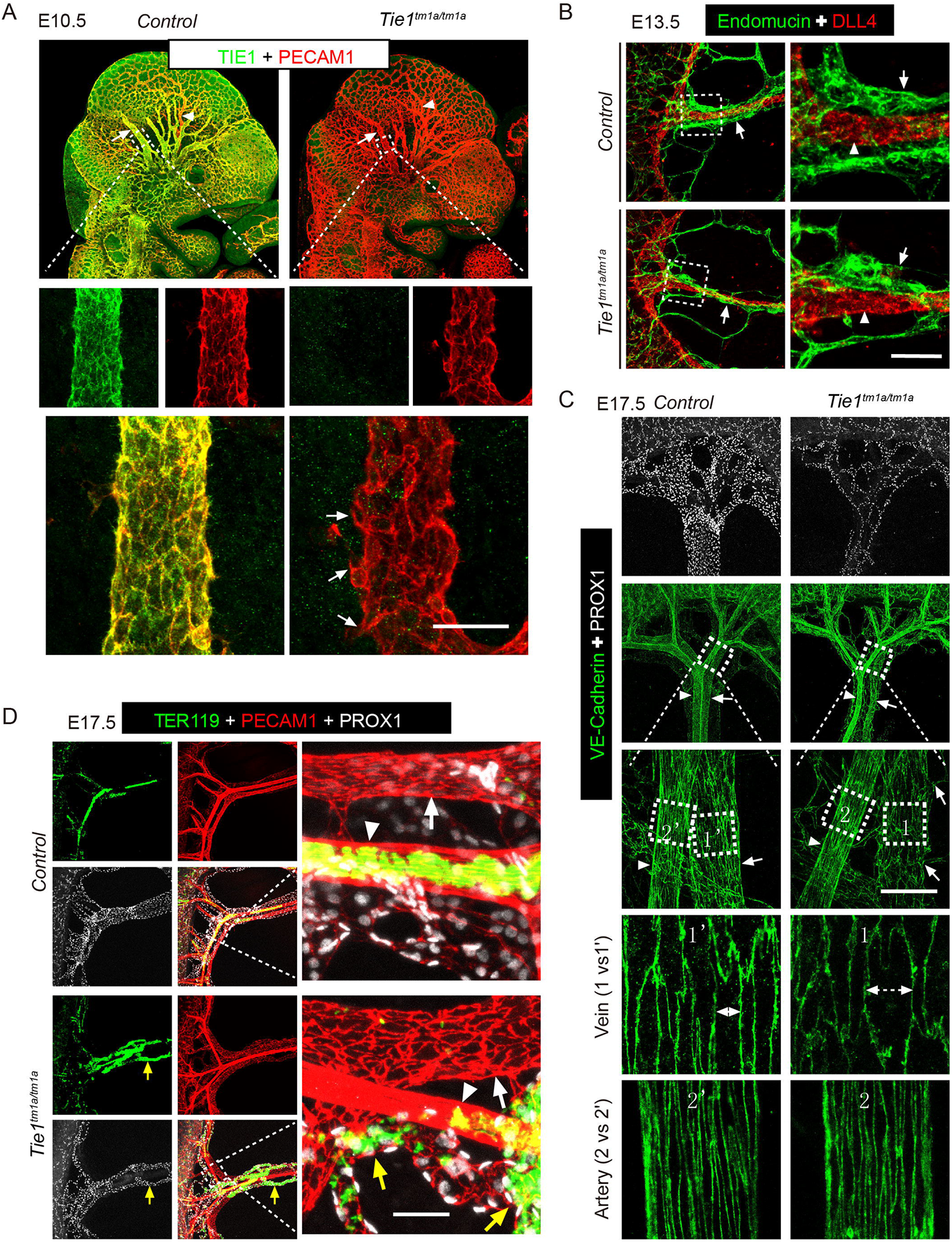
Disruption of venous integrity accompanied by angiogenic sprouting upon *Tie1* deletion. **A**. Disorganized endothelial junctions were observed in the vessel wall of veins in head regions as shown by PECAM1 staining (A, E10.5, *Tie1*^*tm1a/tm1a*^; TIE1, green; PECAM1, red). **B**. The mesentery vein formation was retarded in *Tie1*^*tm1a/tm1a*^ mice compared with that of the littermate controls (E13.5). Arrows point to veins (Endomucin, green) and arrowheads to arteries (DLL4, red). **C**. Analysis of veins and arteries of mesentery (E17.5) as demonstrated by the staining for VE-Cadherin (green). The EC junctions appeared to have an increased distance in the vein walls of *Tie1* mutant mice compared with the controls. Veins were not properly aligned with arteries in the mesentery of *Tie1*^*tm1a/tm1a*^ mice and arrows point to vein-associated angiogenic sprouts in *Tie1* mutant mice (C). Consistent with previous findings, the formation of collecting lymphatics were disrupted in *Tie1* mutant mice (PROX1, white). **D**. Presence of red blood cells in the mesentery lymphatic vessels of *Tie1*^*tm1a/tm1a*^ mice (E17.5, yellow arrows). White arrows point to veins and arrowheads to arteris. Scale bar: 50 μm in A-D.

Consistent with the lymphatic defects observed in *Tie1*^Δ*ICD/*Δ*ICD*^ mutants [12], collecting vessel formation was also disrupted in *Tie1*^*tm1a/tm1a*^ mice compared with the control at E17.5 (Fig. 4C). The loss of vascular integrity was confirmed by the presence of red blood cells detected in the mesenteric lymphatic vessels of *Tie1*^*tm1a/tm1a*^ mice (Fig. 4D, and Supplemental Fig. 3E).

### Alteration of venous and angiogenic gene expression after *Tie1* deletion

RNA sequencing analysis of skin tissues from TIE1 deficient mice (E17.5, *Tie1*^*tm1a/tm1a*^) revealed the altered expression of vascular genes involved in the regulation of venogenesis, angiogenesis as well as hypoxia (Figure 5A-C). The hallmark gene sets, including the hallmarks for vein, artery and angiogenesis used in the GSEA (Gene Set Enrichment Analysis, Supplemental Table 1), are mainly defined according to the transcriptome atlas of murine endothelial cells by Kalucka et al. [37]. Briefly, the top 50 marker genes of venous, angiogenic or arterial endothelial cells identified in ten tissues by Kalucka et al were pooled and the genes expressed in three or more tissues were included in the hallmark gene sets of vein, artery and angiogenesis. The hallmark gene sets of hypoxia and other biological processes are based on the molecular signatures database (h.all.v7.4.symbols) [32, 33]. As shown in Fig. 5B, the venous genes were significantly downregulated whereas arterial genes were not significantly altered in the skin of *Tie1* mutant mice (E17.5). Interestingly, *Tie1* deletion led also to a significant upregulation of genes involved in angiogenesis and hypoxia (Fig. 5B). Heatmaps of the subset of the differentially expressed vein or angiogenesis related genes were shown in Fig. 5C. It is worth pointing out that *Tie1* deletion led to a decrease of venous regulators at the transcript level such as *Tek* and *Aplnr* (Fig. 5C). This was confirmed by the quantitative PCR analysis of skin and lung tissues from *Tie1*^Δ*ICD/*Δ*ICD*^ mice (Fig. 5D). Consistent with the immunofluorescent staining, *Dll4* transcrippts were upregulated while there were no obvious changes with the expression of arterial markers such as *Efnb2* and *Notch1* (Fig. 5D).

**Fig. 5.**
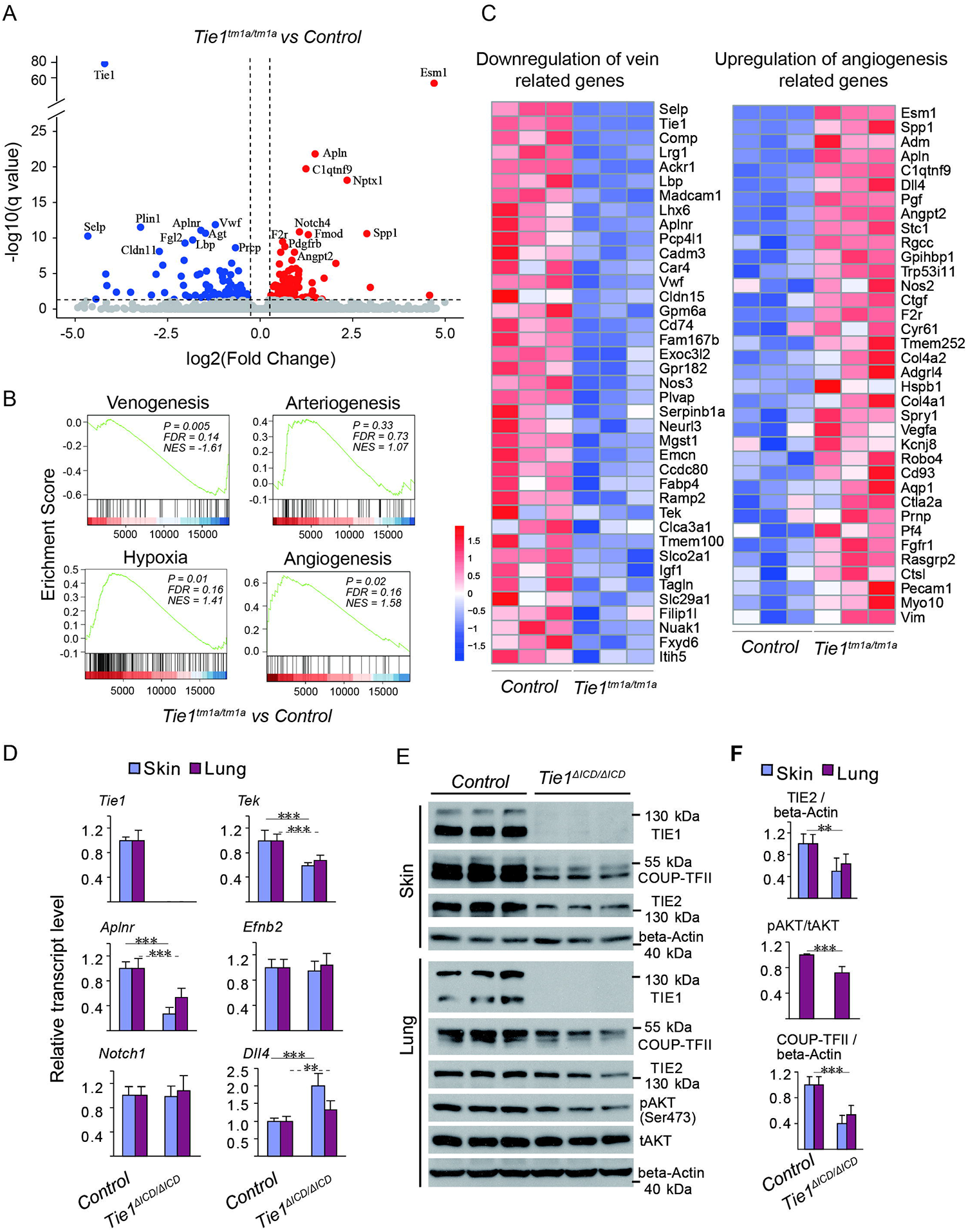
Alteration of venous and angiogenic gene expression after TIE1 deficiency. **A-C**. RNAseq analysis of the skin tissues of *Tie1* null (*Tie1*^*tm1a/tm1a*^) and control mice. The Volcano plot of the differentially regulated genes was shown in A. GSEA analysis revealed that there was a significant decrease in the expression of venous genes but an increase in angiogenesis and hypoxia genes after *Tie1* deletion. In contrast, there was no significant difference in the expression of artery genes (B). The subset of the upregulated angiogenesis genes and down-regulated venous genes (P < 0.05) was shown as heatmaps in C. **D**. Quantitative expression analysis of venous genes including *Aplnr* and *Tek* and arterial genes including *Efnb2, Notch1* and *Dll4* in skin and lung tissues of *Tie1* mutant (*Tie1*^Δ*ICD/*Δ*ICD*^) and control mice (E15.5; Table 1). **E, F**. Western blotting analysis and quantification of COUP-TFII and TIE2 protein normalized against beta-actin in lungs (COUP-TFII/beta-actin: *Tie1*^Δ*ICD/*Δ*ICD*^: 0.56 ± 0.13, n = 6; Control: 1.00 ± 0.11, n = 6. TIE2/beta-actin:*Tie1*^Δ*ICD/*Δ*ICD*^: 0.62 ± 0.17, n = 6; Control: 1.00 ± 0.17, n = 6), and skin of *Tie1* mutant and control mice (COUP-TFII/beta-actin: *Tie1*^Δ*ICD/*Δ*ICD*^: 0.40 ± 0.13, n = 6; Control: 1.00 ± 0.13, n = 6. TIE2/beta-actin:*Tie1*^Δ*ICD/*Δ*ICD*^: 0.50 ± 0.24, n = 6; Control: 1.00 ± 0.18, n = 6). Quantification of AKT (Ser473) phosphorylation in lung is as follows (pAKT / tAKT, *Tie1*^Δ*ICD/*Δ*ICD*^: 0.70 ± 0.21, n=6; Control: 1.00 ± 0.02, n=6).

**Table 1.** The mRNA expression level of venous and arterial endothelial cell markers in skin and lung tissues of *Tie1* knockout and control mice.

| Venous and arterial markers | mRNA expression level (skin) |  | n<br>(*Tie1*<sup>ΔIC</sup><sub>D/ΔICD</sub>) | n<br>(Control) | p value |
| --- | --- | --- | --- | --- | --- |
|  | Control | *Tie1*<sup>ΔICD/ΔICD</sup> |  |  |  |
| *Tie1* | $1.00 \pm 0.06$ | 0 | 9 | 10 | <0.0001 |
| *Tek* | $1.00 \pm 0.16$ | $0.59 \pm 0.05$ | 9 | 10 | <0.0001 |
| *Aplnr* | $1.00 \pm 0.10$ | $0.27 \pm 0.10$ | 9 | 10 | <0.0001 |
| *Efnb2* | $1.00 \pm 0.13$ | $0.95 \pm 0.15$ | 9 | 10 | 0.454 |
| *Notch1* | $1.00 \pm 0.14$ | $0.98 \pm 0.17$ | 9 | 10 | 0.733 |
| *Dll4* | $1.00 \pm 0.07$ | $2.00 \pm 0.36$ | 9 | 10 | <0.0001 |

Lung transcript levels of venous and arterial markers
| Venous and arterial markers | mRNA expression level (lung) |  | n<br>(*Tie1*<sup>ΔIC</sup><sub>D/ΔICD</sub>) | n<br>(Control) | p value |
| --- | --- | --- | --- | --- | --- |
|  | Control | *Tie1*<sup>ΔICD/ΔICD</sup> |  |  |  |
| *Tie1* | $1.00 \pm 0.16$ | 0 | 9 | 9 | <0.0001 |
| *Tek* | $1.00 \pm 0.10$ | $0.67 \pm 0.1$ | 9 | 9 | <0.0001 |
| *Aplnr* | $1.00 \pm 0.16$ | $0.53 \pm 0.15$ | 9 | 9 | <0.0001 |
| *Efnb2* | $1.00 \pm 0.14$ | $1.04 \pm 0.19$ | 9 | 9 | 0.618 |
| *Notch1* | $1.00 \pm 0.14$ | $1.07 \pm 0.25$ | 9 | 9 | 0.458 |
| *Dll4* | $1.00 \pm 0.13$ | $1.31 \pm 0.28$ | 9 | 9 | 0.00684 |

Consistent with the downregulation of *Tek* expression, *Tie1* deletion led to a decrease of TIE2 as well as COUP-TFII proteins as demonstrated in the skin and lung tissues of *Tie1*^Δ*ICD/*Δ*ICD*^ mutant mice (Figure 5E and 5F). There was also a significant decrease of AKT phosphorylation in the lung of *Tie1* mutants (Ser473, Figure 5F), which may account for the decrease of COUP-TFII as demonstrated in *Tek* deficient mice [3]. Furthermore, decrease of TIE2 and COUP-TFII proteins was further verified in cultured human umbilical vein endothelial cells (HUVECs) when TIE1 was reduced by the siRNA mediated knockdown (Figure 6A-C). This suggests that TIE1 may function in vein development, at least partly, by regulating the expression of TIE2.

**Figure 6.**
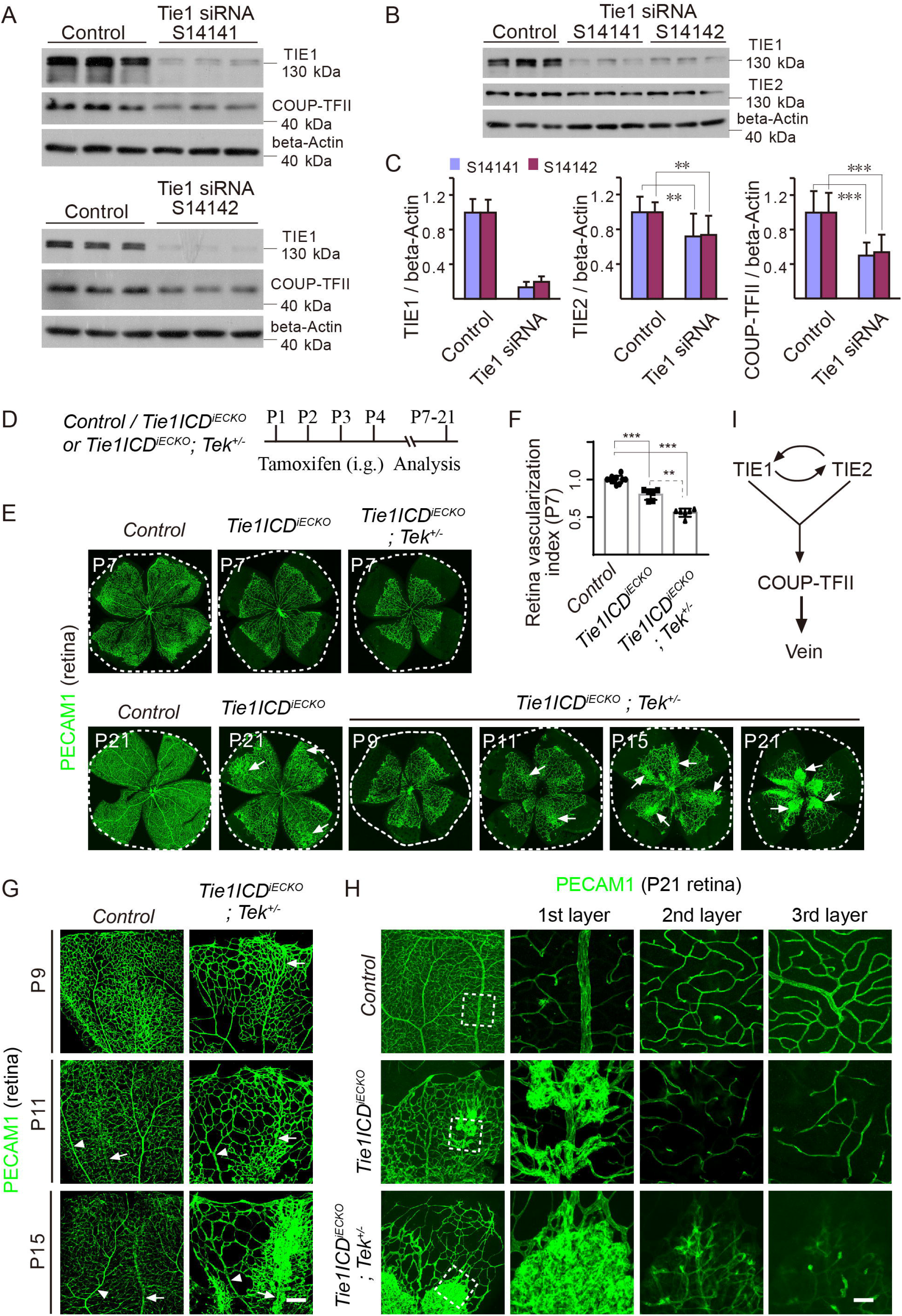
Synergy of TIE1 and TIE2 in retinal venous development. **A-C**. Western blot analysis of COUP-TFII protein in HUVECs after siRNA mediated *Tie1* knockdown. Two small interfering (siRNAs) targeting *Tie1* were used in this study, including S14141 and S14142. Quantification of TIE1, TIE2 and COUP-TFII protein level was as follows. For S14141: COUP-TFII was decreased to 61.44 ± 27.32 % and TIE2 to 71.63 ± 25.52 % of the control respectively, when TIE1 expression was reduced to 15.11 ± 6.85 % of control (normalized by beta-actin; values from five independent experiments, n=15). For S14142: COUP-TFII was decreased to 53.92 ± 20.29 % and TIE2 to 72.39 ± 24.06 % of the control respectively, when TIE1 expression was reduced to 20.30 ± 6.12 % of control (normalized by beta-actin; values from three independent experiments, n=9). **D**. Tamoxifen intragastric (i.g.) administration and the analysis scheme. **E, F**. Analysis of blood vessels in the retinas of *Tie1ICD*^*ECKO/-*^, *Tie1ICD*^*ECKO/-*^*;Tek*^*+/-*^ and control mice between P7 and P21. Arrows point to haemangioma-like vascular tufts (E). Quantification of the vascularization index (F; ratio of vascularized area to total retina area normalized against the littermate controls at P7; Control: 100.00 ± 5.06, n=10; *Tie1ICD*^*ECKO/-*^: 80.00 ± 7.03, n=6; *Tie1ICD*^*ECKO/-*^; *Tek* ^*+/-*^: 55.92 ± 5.42, n=6; Control versus *Tie1ICD*^*ECKO/-*^: *P*=0.0001; Control versus *Tie1ICD*^*ECKO/-*^*;Tek*^*+/-*^: *P*=0.0001; *Tie1ICD*^*ECKO/-*^ versus *Tie1ICD*^*ECKO/-*^*;Tek*^*+/-*^: *P*=0.0022). **G-H**. Visualization of retinal blood vessels of *Tie1ICD*^*ECKO/-*^*;Tek*^*+/-*^ mice by immunostaining for PECAM-1 at P9, P11, P15 and P21. Arrows point to veins and arrowheads to arteries (G). Analysis of the three layers of retinal blood vessels in *Tie1ICD*^*ECKO/-*^, *Tie1ICD*^*ECKO/-*^*;Tek*^*+/-*^ and control mice at P21 (H). Note that the vein-associated vascular defects were more severe in *Tie1ICD*^*ECKO/-*^*;Tek*^*+/-*^ mice compared with the *Tie1* deletion alone. **I**. Schematic model shows that TIE1 and TIE2 act in a synergistic manner to regulate vein development via COUP-TFII. Scale bar: 200 μm in G; 50 μm in H.

### Synergy of TIE1 and TIE2 in retinal vein morphogenesis

To further confirm the synergistic effect of TIE1 and TIE2 on vascular development, particularly vein formation, we generated mice with the EC-specific deletion of *Tie1* (*Tie1ICD*^*iECKO*^: *Tie1ICD*^*Flox/-*^*/Cdh5-Cre*^*ERT2*^) [12, 25], and also the compound knockout mice targeting *Tie1* plus one null allele of *Tek* (*Tie1ICD*^*iECKO*^*;Tek*^*+/-*^). The scheme for *Tie1* deletion by the intragastric administration of tamoxifen is shown in Fig. 6D. Deletion of *Tie1* alone (*Tie1ICD*^*iECKO*^, P1-4) led to a significant decrease in the retinal vascularization at the postnatal day 7 (P7; Fig. 6E-F). Interestingly, lack of TIE1 plus one null allele of *Tek* (*Tie1ICD*^*iECKO*^*;Tek*^*+/-*^) resulted in a more severe decrease in the retinal blood vessel growth (Fig. 6E-F). Quantification of the retinal vascularization index was performed as previously published [3], and shown in Fig. 6F (Control: 100.00 ± 5.06, n=10; *Tie1ICD*^*iECKO*^: 80.00 ± 7.03, n=6; *Tie1ICD*^*iECKO*^*;Tek*^*+/-*^: 55.92 ± 5.42, n=6; Control versus *Tie1ICD*^*iECKO*^: *P*=0.0001; Control versus *Tie1ICD*^*iECKO*^*;Tek*^*+/-*^: *P*=0.0001; *Tie1ICD*^*iECKO*^ versus *Tie1ICD*^*iECKO*^*;Tek*^*+/-*^: *P*=0.0022) (Figure 6F).

In addition, similar to the vascular defects observed in the retinas of *Tek* deleted mice [3], there was also a progressive increase of vein-associated angiogenesis leading to the formation of haemangioma-like vascular tufts in *Tie1ICD*^*iECKO*^*;Tek*^*+/-*^ mice (from P9 to P21; Fig. 6E, G, H and Supplemental Fig. 5A-C). The retinal veins and arteries were identified by the immunostaining for EphB4 and DLL4 (Supplemental Fig. 5B-C). Vascular growth towards the deep layers of retinas was diminished in *Tie1ICD*^*iECKO*^ and *Tie1ICD*^*iECKO*^*;Tek*^*+/-*^ mice compared with the littermate controls (Fig. 6H). Notably, the retinal venous defects were more severe in *Tie1ICD*^*iECKO*^*;Tek*^*+/-*^ mice than the mutant mice with *Tie1* deletion alone (*Tie1ICD*^*iECKO*^; Fig. 6E, H), suggesting that TIE1 and TIE2 function in a synergistic manner in the regulation of retinal vascularization, particularly in retinal vein development (Fig. 6I).

## Discussion

Functional mechanisms of endothelial TIE receptors have attracted intensive research over the last three decades since their discovery. The goal of this study is to investigate roles of TIE1 and its relationship with TIE2 in the establishment of veins. We show evidence in this study that TIE1 plays a crucial role in restricting angiogenesis for the assembly of veins and acts in a synergistic manner with TIE2 in this process.

TIE1 and TIE2 are differentially required during the blood vascular and lymphatic network formation [9, 15, 38]. Recent studies have shown that TIE1 plays important roles in the regulation of blood vascular growth [16-18], in addition to its requirement in the formation of collecting lymphatics [12, 15, 36, 39]. TIE2 is required for the formation and maturation of blood vascular network, especially in vein development [3, 9]. We show in this study that the formation of cardinal veins is independent of TIE1. Cardinal veins are formed by a process of vasculogenesis, involving *de novo* formation of vessels from angioblasts [40]. This suggests that TIE1 is not essential for the differentiation of venous ECs. Consistent with the previous observation [3], TIE2 is critical in the specification of cardinal venous ECs as evidenced by the aberrant expression of DLL4 upon the loss of TIE2. Interestingly, in contrast to the complete lack of cutaneous veins in *Tek* mutant mice [3], lack of TIE1 disrupted the initial formation of veins in skin. Interestingly, cutaneous veins were detected at later stages of embryonic development but displaying the arteriovenous misalignment. Therefore, TIE1 deficiency delayed the formation of veins, suggesting the differential requirement of TIE1 and TIE2 during the vein development.

In spite of the distinct functions, we have also found that TIE1 and TIE2 act in a synergistic manner in the vein development. Induced deletion of *Tek* in neonatal mice led to the formation of angioma-like vascular tufts resulting from uncontrolled vein-associated angiogenesis in retinas [3]. Loss of TIE1 resulted in a similar but relatively mild increase of retinal vein-associated angiogenesis in comparison with that of *Tek* mutants [3]. Although there was no obvious vascular abnormality with mice heterozygous for *Tek* deletion in retinas, combining the induced endothelial knockout of *Tie1* with one null allele of *Tek* produced a more severe vein-associated vascular phenotype in retinas than that of *Tie1* deletion alone, with a comparable formation of the vascular tufts as those of *Tek* mutant mice. This points to a synergistic function of TIE1 and TIE2 in the restriction of sprouting angiogenesis during the vein formation. Synergistic roles of TIE1 and TIE2 in blood vascular formation were also demonstrated in other studies. Insufficient TIE1 decreased the ANGPT1-induced TIE2 activation and suppressed the enlargement of tracheal vessels by the treatment with angiopoietins [16]. Delivery of soluble TIE2 by the adeno-associated virus (AAV) vectors plus the genetic deletion of *Tie1* produced an additive effect on the inhibition of tumor angiogenesis [18]. Furthermore, both TIE1 and TIE2 could regulate the expression of COUP-TFII, a key factor for the venous endothelial cell fate. Consistent with the findings in *Tek* mutant mice [3], we observed that COUP-TFII expression decreased in mice by the genetic deletion of *Tie1* or in cultured endothelial cells by siRNA-mediated knockdown of *Tie1*. TIE1 deficiency also disrupted the arterial-venous alignment in tissues including skin and mesentery. This resembles the abnormal arteriovenous alignment of the *Aplnr* (also known as APJ) or *Tek* deficient mice [3, 41]. A significant reduction of *Aplnr* expression was detected in the *Tie1* deficient mice, which was also observed in *Tek* mutants as previously published [3]. As the TIE1 deficiency reduced the expression of *Tek*, it is likely that TIE1 exerts its function, at least partly, via the regulation of TIE2 in the venous development.

How do TIE receptors exert their synergistic functions during the vascular development? TIE2 activation suppresses the Forkhead box protein O1 (FOXO1)-mediated transcriptional regulation via AKT pathway and inhibition of TIE2 activates FOXO1, leading to the increase of ANGPT2 expression [23, 42]. Induced deletion of *Tie1* has also been shown to increase FOXO1 nuclear localization and transcriptional activation [16]. In this study, we found that *Tie1* deletion led to a decrease in the expression of venous endothelial related genes in embryonic skin such as *Tek, Aplnr, Emcn*, and an increase of angiogenic regulators including *Angpt2, Vegfa* and *Dll4*. This may account for the vein-associated increase of angiogenesis during embryogenesis and also in the retinas after the postnatal deletion of *Tie1*, which became more severe in the double mutants combining the endothelial *Tie1* deletion with one null allele of *Tek*.

Furthermore, loss of TIE1 led to the disruption of vascular integrity as demonstrated by the vascular bleeding. Platelets play a central role in primary hemostasis during physiological and pathological conditions [43]. It is worth noting that TIE1 deficiency leads to the decrease of P-selectin expression, which may interfere with the endothelial-platelet interaction [44]. In addition, the decrease of VWF expression upon TIE1 deficiency also points to a role of TIE1 in the regulation of the coagulation system for vascular integrity. Besides the above discussed, endothelial adherens junctions are among the key regulatory components for the establishment and homeostasis of endothelial integrity [45]. TIE1 and TIE2 are expressed by endothelial tip and/or stalk cells [17]. In cultured endothelial cells, TIE1 and TIE2 are translocated to endothelial cell contacts upon ANGPT1 treatment [46], suggesting that TIE receptors are important components of the intercellular adhesions in addition to transducing signals for the assembly of vascular wall.

In summary, findings from this study imply that TIE1 and TIE2 act as key regulators of multiple events during the vein formation, including the restriction of sprouting angiogenesis, specification of venous endothelial cells, and regulation of endothelial cell junctions for vessel wall integrity.

## Acknowledgement

We thank the staff in Animal facility of Soochow University for technical assistance.

## Highlights

1. TIE1 was dispensable for cardinal vein development while TIE2 deficiency altered the identity of cardinal vein endothelial cells with the aberrant expression of DLL4.
2. Loss of TIE1 disrupted the vein formation in skin during embryogenesis as well as in the retinas after the postnatal endothelial *Tie1* deletion.
3. TIE1 deficiency led to the decreased expression of venous regulators including TIE2 while angiogenic regulators were upregulated.
4. Combining the endothelial deletion of *Tie1* with one null allele of *Tek* resulted in a more severe retinal vein defects than that of *Tie1* deletion alone, suggesting that TIE1 and TIE2 act in a synergistic manner in venogenesis.

