## Supplemental files for "Endothelial TIE1 restricts angiogenic sprouting to coordinate vein assembly in synergy with its homologue TIE2"

### Supplemental materials
### Supplemental figures and figure legends

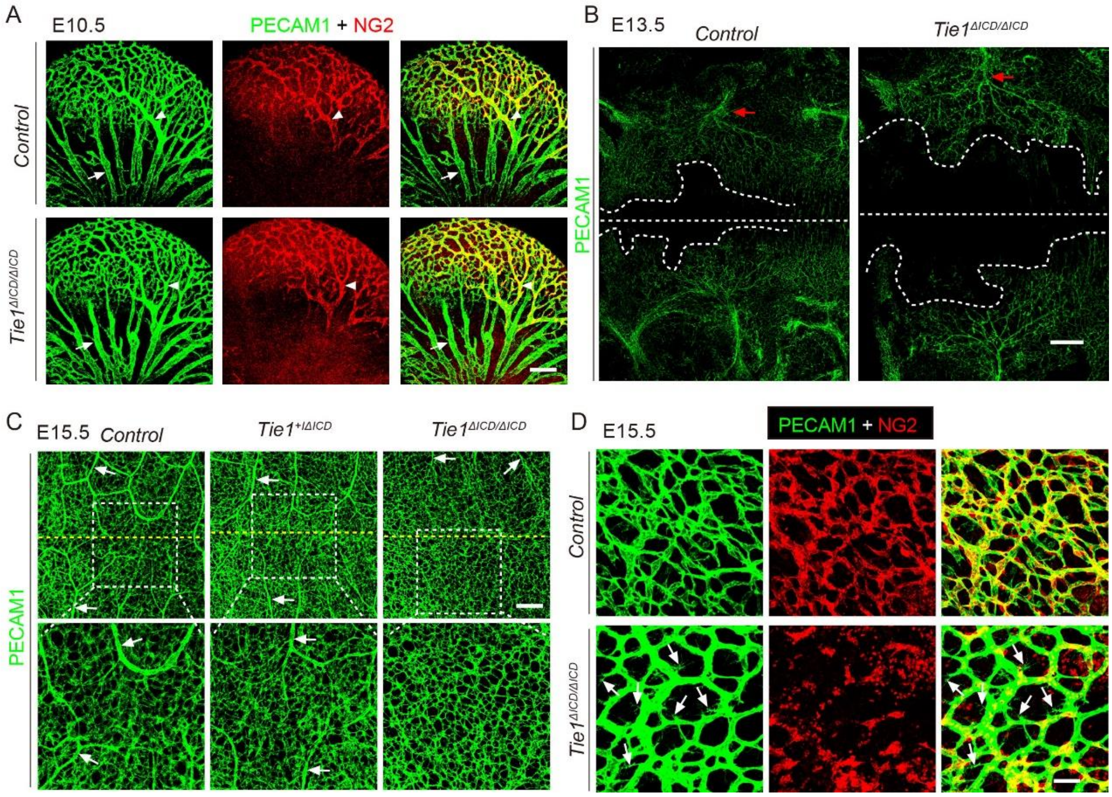


**Supplemental Fig. 1 TIE1 deficiency retarded the vein formation and vascular growth toward the middle line regions of dorsal skin. A**. Analysis of vein formation and vascular mural cells in head regions of *Tie1*$^{\Delta ICD/\Delta ICD}$ mice (E10.5) by immunostaining for PECAM1 (green) and NG2 (red). Arrows point to veins and arrowheads to arteries. **B**. Vascular growth (PECAM1, green) toward the middle line of the skin was delayed in the *Tie1* mutant mice compared with the littermate controls (E13.5). Note that the curved dotted lines indicate the non-vascularized regions in the back skin of *Tie1* mutant and control mice. **C**. Arteries and veins converged at middle lines of back skin in wildtype control mice, while veins were not properly formed in *Tie1* mutant mice (C, E15.5). Arrows point to small arteries (strong PECAM1 staining) approaching the middle-line regions of back skin (E15.5). **D**. In comparison with the well-formed blood vascular network in the control mice (E15.5), there was still active angiogenesis (arrows point to the angiogenic sprouts) in the skin of *Tie1* mutants, which was also shown to have less pericyte coverage (NG2, red; PECAM1, green). Straight dotted lines indicate the middle line regions of the back skin. Scale bar: 200 μm in A, C; 500 μm in B; 50 μm in D.

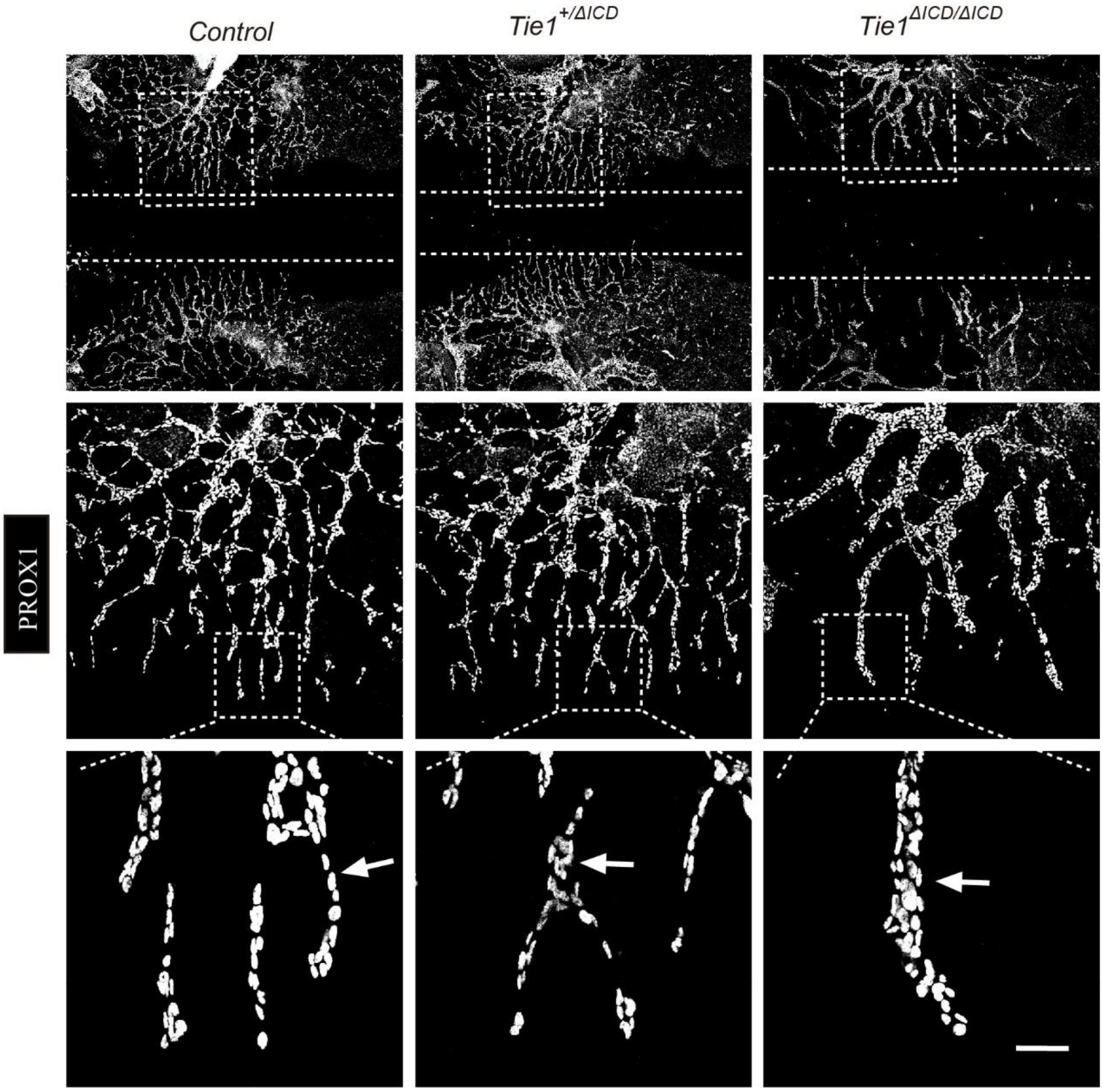


**Supplemental Fig. 2 Retardation of lymphatic vessel growth in the skin of *Tie1*$^{\Delta ICD/\Delta ICD}$ mice.** Analysis of lymphatic vessel formation by immunostaining for PROX1 (white) with the back skin of embryos (E13.5). Note the lymphatic vessel growth toward the middle line of back skin (dotted lines) was retarded and lymphatic vessels became dilated in the *Tie1*$^{\Delta ICD/\Delta ICD}$ mutants compared with those of control mice (arrows). Scale bar: 50 μm.

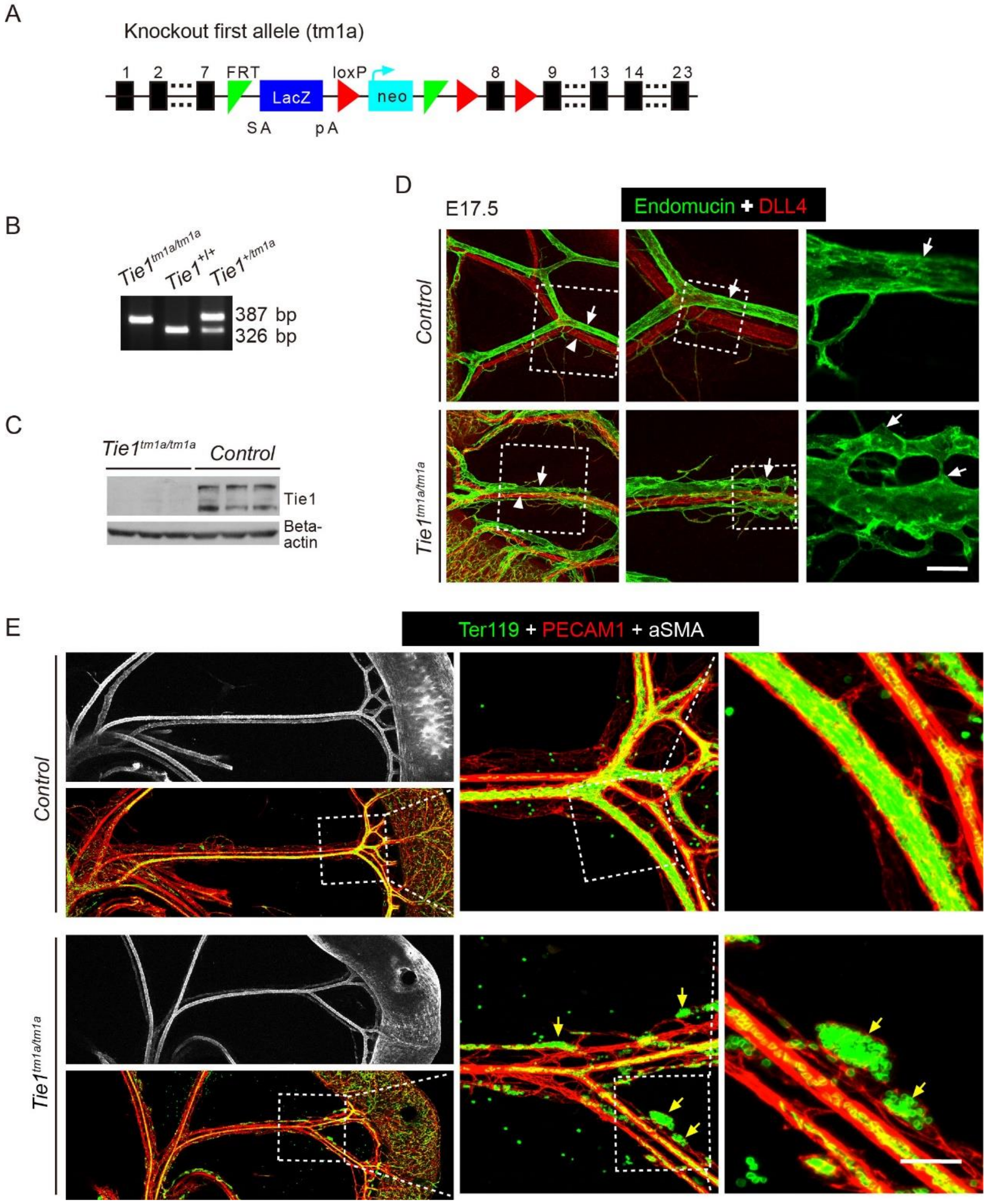


**Supplemental Fig. 3 Generation of a conditional knockout mouse line targeting *Tie1* and analysis of the mesentery vein formation during embryogenesis. A-C**. Targeting strategy for the *Tie1* knockout first allele (A, *Tie1*$^{tm1a}$), the genotyping (B) and western blotting analysis of TIE1 (C) in *Tie1*$^{tm1a/tm1a}$ and control mice. **D**. Analysis of the mesentery vein formation in *Tie1*$^{tm1a/tm1a}$ and control mice at E17.5. Note that the active angiogenic sprouting was still ongoing in veins of *Tie1*$^{tm1a/tm1a}$ mice compared with the well-formed veins in the littermate controls. Arrows point to veins (Endomucin, green) and arrowheads to arteries (DLL4, red). **E**. Analysis of vascular bleeding in the mesentery of *Tie1*$^{tm1a/tm1a}$ mice at E17.5. Note that there were red blood cells outside of blood vessels in the mesentery of *Tie1* mutant mice. Scale bar: 50 μm in D and E.

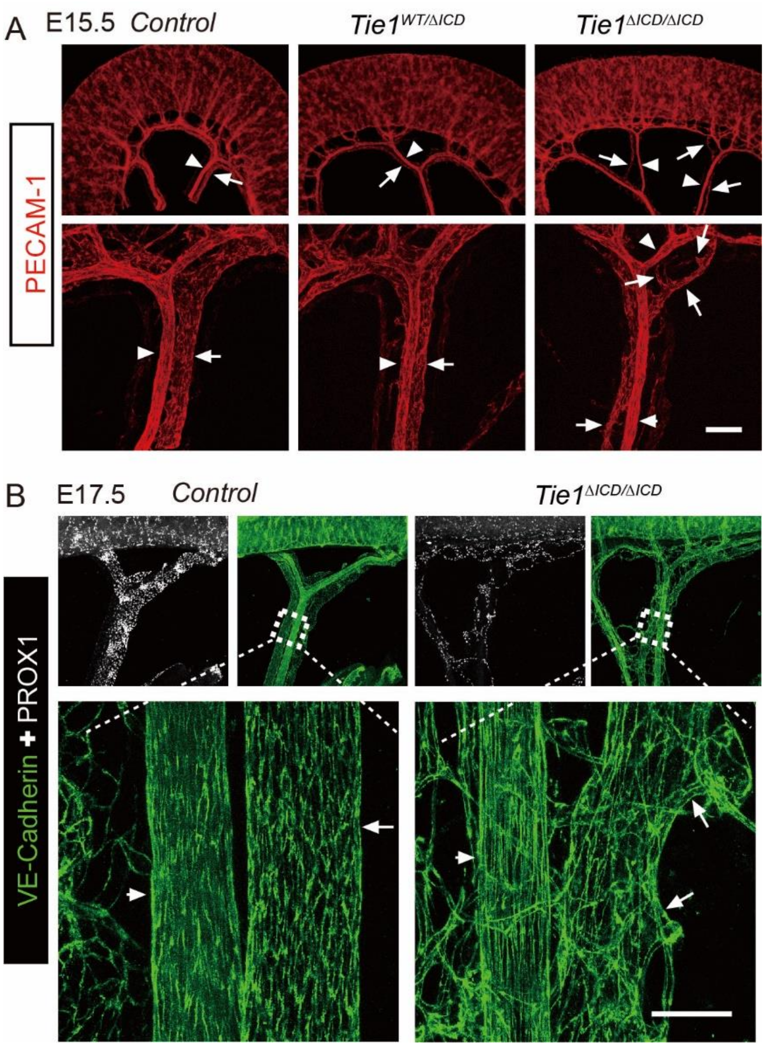


**Supplemental Fig. 4 Vein-associated sprouting angiogenesis and misalignment of veins with arteries in the mesentery of *Tie1*$^{\Delta ICD/\Delta ICD}$ mice. A-B**. Misalignment of veins with arteries, together with vein-associated angiogenic sprouts, were also detected in the mesentery of *Tie1*$^{\Delta ICD/\Delta ICD}$ mice (A, E15.5; B, E17.5). Scale bar: 50 μm in A and B.

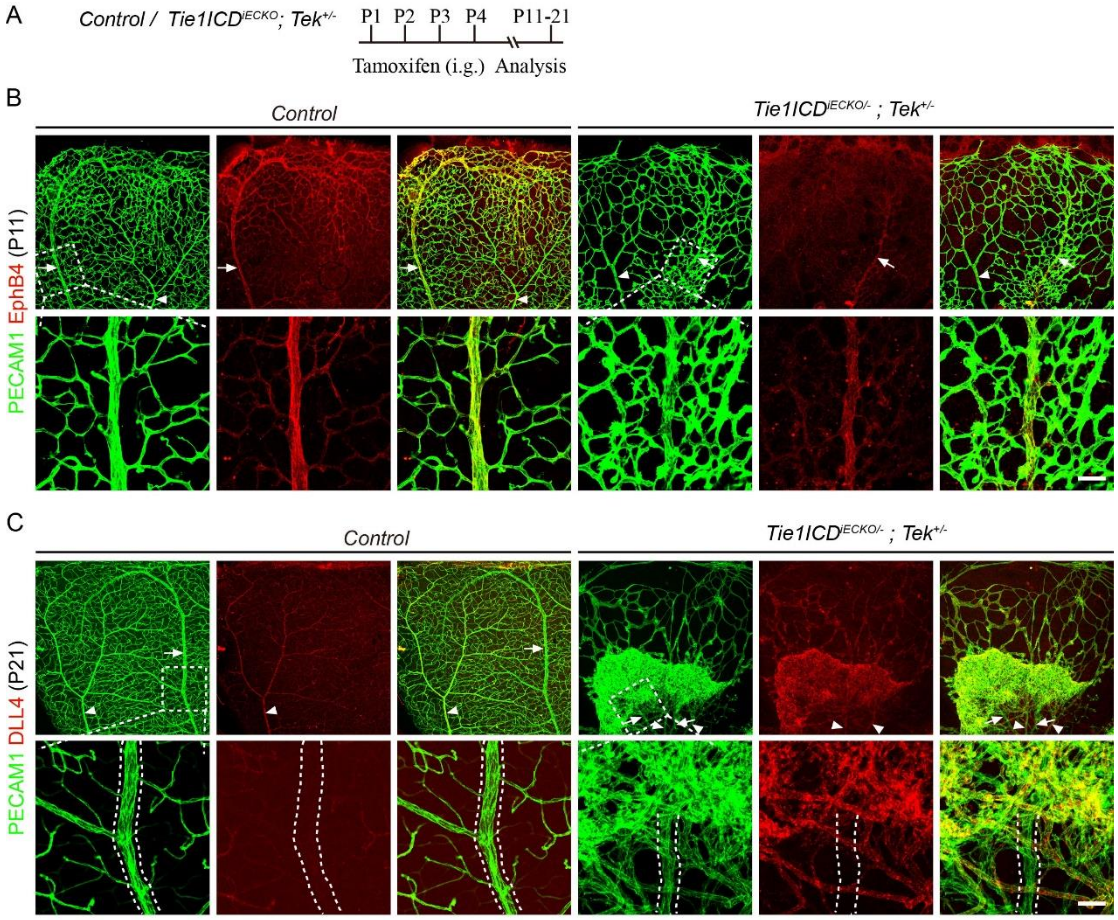


**Supplemental Fig. 5 Combining the endothelial *Tie1* deletion plus one null allele of *Tek* leads to a dramatic increase of vein-associated angiogenesis in retina. A.** Tamoxifen intragastric (i.g.) administration and the analysis scheme. **B,C**. Analysis of blood vessels in the retinas of *Tie1ICD*$^{iECKO/-}$;*Tek*$^{+/-}$ mice and control mice at P11 (B), and P21 (C). Arrows point to veins (EphB4 positive in B and DLL4 negative in C) and arrowheads to arteries (DLL4 positive in C). Note that retinas in *Tie1ICD*$^{iECKO/-}$;*Tek*$^{+/-}$ mice showed a dramatic increase of vein-associated angiogenesis leading to the formation of vascular tufts at P21. Scale bar: 50 μm in B and C.

**Supplemental table 1**

| **Supplemental Table 1 Hallmark gene sets of vein, artery and angiogenesis used for the GSEA analysis** | |
|---|---|
| HALLMARK_VEIN | Abcg2, Ackr1, Car8, Cfh, Cytl1, Dnm3, Emcn, Fmo2, Gm13889, Id2, Lhx6, Mgp, Pdzd2, Pltp, Prpf40b, Tshz2, Ctsh, Ier3, Nfkbia, Nr2f2, Nuak1, Pdia5, Selp, Slco2a1, Vim, 2200002D01Rik, Amigo2, AU021092, Calcrl, Cpe, Eln, Klk8, Prss23, Rasa4, Rbp1, Spint2, Tmem252, Ehd4, Lrg1, Plvap, Tmem176a, Hs3st1, Tmem176b, Apoe, Ctla2a, Icam1, Il6st, Ptgs1, Tmsb10, Vcam1, Bgn, Vwf, Tek, Tie1, Aplnr, Ephb4, Pik3ca, Nos3, Angpt1 |
| HALLMARK_ARTERY | Bmx, Bsg, Cd9, Eln, Eps8l2, Hist1h2bc, Hspa1a, Ifi27l2a, Jund, Ly6a, Nr4a1, Slc3a2, Tinagl1, Tppp3, Tspan7, Vim, Zfp36, Adamts1, Cdh13, Dll4, Ebf1, Fn1, Fos, Junb, Ltbp4, Nrarp, Rbp7, Ssu2, Alpl, Amd1, Cyr61, Htra1, Jag1, Klf4, S100a6, Ace, Dusp1, Gadd45g, Gja5, Glul, Ier2, Igfbp3, Mgp, Plat, Cst3, Edn1, Fbln5, Id1, Igfbp4, Ptprr, Slc6a6, Vegfc, 8430408G22Rik, Azin1, Fbln2, Gja4, Hey1, Mecom, Sat1, Sox17, Stmn2, Tsc22d1, Clu, Sema3g, Tm4sf1, Crip1, Efnb2, Notch1, Nrp1 |
| HALLMARK_ANGIOGENESIS | Adm, Apln, Col15a1, Col4a1, Col4a2, Trp53i11, Actb, Bcl6b, Ccnd1, Gpihbp1, Lamc1, Lgals1, Lpar6, Sox4, Sparc, Vwa1, Angpt2, Gm12002, Itga6, Noct, Scarb1, Spry1, Tubb6, Adora2a, Arl4c, Bcl2, Cd247, Gadd45a, Git2, Il16, Msmp, Mycn, Pcdh12, Rnf144a, Tnfaip8l1, Kdr, Vegfa, Hif1a, Esm1, Dll4, Flt1 |
